# Total whole-arm chromosome losses predict malignancy in human cancer

**DOI:** 10.1101/2025.03.09.642243

**Authors:** Ye Zheng, Kami Ahmad, Steven Henikoff

## Abstract

Aneuploidy is observed as gains or losses of whole chromosomes or chromosome arms and is a common hallmark of cancer. Whereas models for the generation of aneuploidy in cancer invoke mitotic chromosome segregation errors, whole-arm losses might occur simply as a result of centromere breakage. We recently showed that elevated RNA Polymerase II (RNAPII) level over S-phase-dependent histone genes predicts rapid recurrence of human meningioma and is correlated with total whole-arm losses relative to gains. To explain this imbalance in arm losses over gains, we have proposed that histone overexpression at S-phase competes with the histone H3 variant CENP-A, resulting in centromere breaks and whole-arm losses. To test whether centromere breaks alone can drive aneuploidy, we ask whether total whole-arm aneuploids can predict outcome across different cancer types in large RNA and whole-genome sequencing databanks. We find that total whole-arm losses generally predict outcome, suggesting that centromere breakage is a major initiating factor leading to aneuploidy and the resulting changes in the selective landscape that drive most cancers. We also present evidence that centromere breakage alone is sufficient to account for whole-arm losses and gains, contrary to mitotic spindle error models for generation of aneuploidy. Our results suggest that therapeutic intervention targeting histone overexpression has the potential of reducing aneuploidy and slowing cancer progression.

**Significance Statement:** Gain or loss of whole chromosome arms following centromere breaks is frequent in cancer, but whether or not there is a common initiating event is unknown. Here we show that the total number of whole-arm losses predicts patient outcomes across cancer types, suggesting a causal relationship. This general excess of losses over gains is not predicted by mitotic error models of aneuploidy but rather suggests that centromere breaks themselves initiate whole-arm aneuploidies. Insofar as aneuploidy reshapes the selective landscapes that drive most cancers, our results have potential clinical implications.

## Introduction

Aneuploidy is a familiar hallmark of cancer that was first described well over a century ago (1). In 1890, David Hansemann observed asymmetric mitoses in a variety of epithelial cancers but not in normal tissues. Among the forms that Hansemann illustrated were examples of chromatids away from the metaphase plate that were either attached or unattached to the mitotic spindle, in addition to many examples of multipolar spindles. More than 130 years after Hansemann’s observations, the causes of aneuploidy have been understood as belonging to any of four classes of mitotic errors (2): merotelic attachments, where a single kinetochore connects to opposite spindles, resulting in a chromosome that remains at the metaphase plate; extra centrosomes, where chromosomes segregate to three or more poles; unattached kinetochores, where only one sister chromatid is attached and both are pulled to the same pole; or cohesion defects where sister chromatids either release from one another prior to anaphase or fail to release at anaphase. Notably, each of these chromosome instability mechanisms can result in the gain or loss of chromosomes or chromosome fragments based on cytological preparations of tumors. This diversity of mechanisms that are thought to drive aneuploidy in cancer severely complicates therapeutic strategies.

In recent years, whole-genome sequencing (WGS) and RNA sequencing (RNA-seq) have provided efficient alternatives to karyotype analysis for scoring aneuploidies in cancer patient samples. Whole-chromosome, whole-arm and partial gains or losses can be accurately scored by measuring differences in DNA or RNA abundances across the genome. These genomic and transcriptomic studies have led to the realization that aneuploidy encompasses multiple varieties of somatic copy number alterations generated by different molecular mechanisms (3). Whole-arm aneuploidy is especially common in cancer and certain losses or gains are important prognostic indicators. For example, in myelodysplastic syndrome, which can lead to acute myeloid leukemia, loss of chromosome arm 5q predicts a more positive outcome, while loss of chromosome 7 or arm 7q predicts a more negative outcome after bone marrow transplantation, and these indicators have long been used to determine the course of treatment (4).

WGS or RNA-seq of routine patient samples is often challenging because clinical samples are typically banked as formalin-fixed paraffin-embedded sections (FFPEs), but we recently implemented RNA Polymerase II (RNAPII) profiling in FFPEs as an efficient method for assessing chromatin profiling in clinical samples (5). We also surveyed a variety of patient tumor and normal samples, including 30 meningiomas and 15 breast tumors. Using the RNAPII signal at candidate cis-regulatory elements (cCREs) from the Encyclopedia of DNA Elements (ENCODE) to identify whole-arm aneuploids, we found that the total number of whole-arm aneuploids accurately predicted rapid recurrence in meningiomas (6). RNAPII at S-phase-dependent histone genes strongly correlated with whole-arm losses relative to gains in both meningiomas and breast tumors, suggesting a causal relationship.

Here we use available public RNA-seq and WGS datasets to test the generality of our predictions based on total arm aneuploid counts using RNAPII data. Our analyses confirm that the total number of whole-arm losses predicts outcome better than gains. We propose that whole-arm losses are immediate consequences of most centromere breaks, and others with partially intact centromeres form micronuclei, where they undergo S-phase replication and reattachment at a subsequent anaphase.

## Results

### Whole-arm losses correlate with recurrence in meningioma RNA-seq and pan-cancer WGS data

Whole-arm losses and gains of metacentric chromosomes are generated by breaks in centromeres or pericentromeric regions, although the underlying mechanisms have been speculative (7). We previously showed that RNAPII at histone genes in 30 patient samples predicted rapid recurrence when integrated with RNA-seq data from 1298 meningiomas. We also showed that RNAPII at histone genes correlated with whole-arm losses relative to whole-arm gains, and we wondered whether the meningioma RNA-seq data shows the same bias of losses over gains. S-phase-dependent histone mRNAs are the only RNAPII-transcribed protein-coding genes that are not 3’-polyadenylated, and so they are grossly under-represented in RNA sequencing datasets (Fig. S1).

To determine whole-arm gains and losses in RNA-sequencing (RNA-seq) data, CaSpER was leveraged to identify the copy number variants (CNVs) and, hence, determine the whole-arm gains or losses according to the consistency-based large-scale CNV calls (8, 9). We observed very low levels of recurrence with <3 whole-arm losses, intermediate levels of recurrence with 3-6 losses and highest levels of rapid recurrence with >6 losses (Fig. 1A, left panel). Low levels of recurrence were also observed for <3 gains, however, >6 gains were not associated with rapid recurrence (Fig. 1A, middle panel). Total whole-arm aneuploidy also predicted recurrence, although not as well as whole-arm losses (Fig. 1A, right panel).

**Figure 1.**
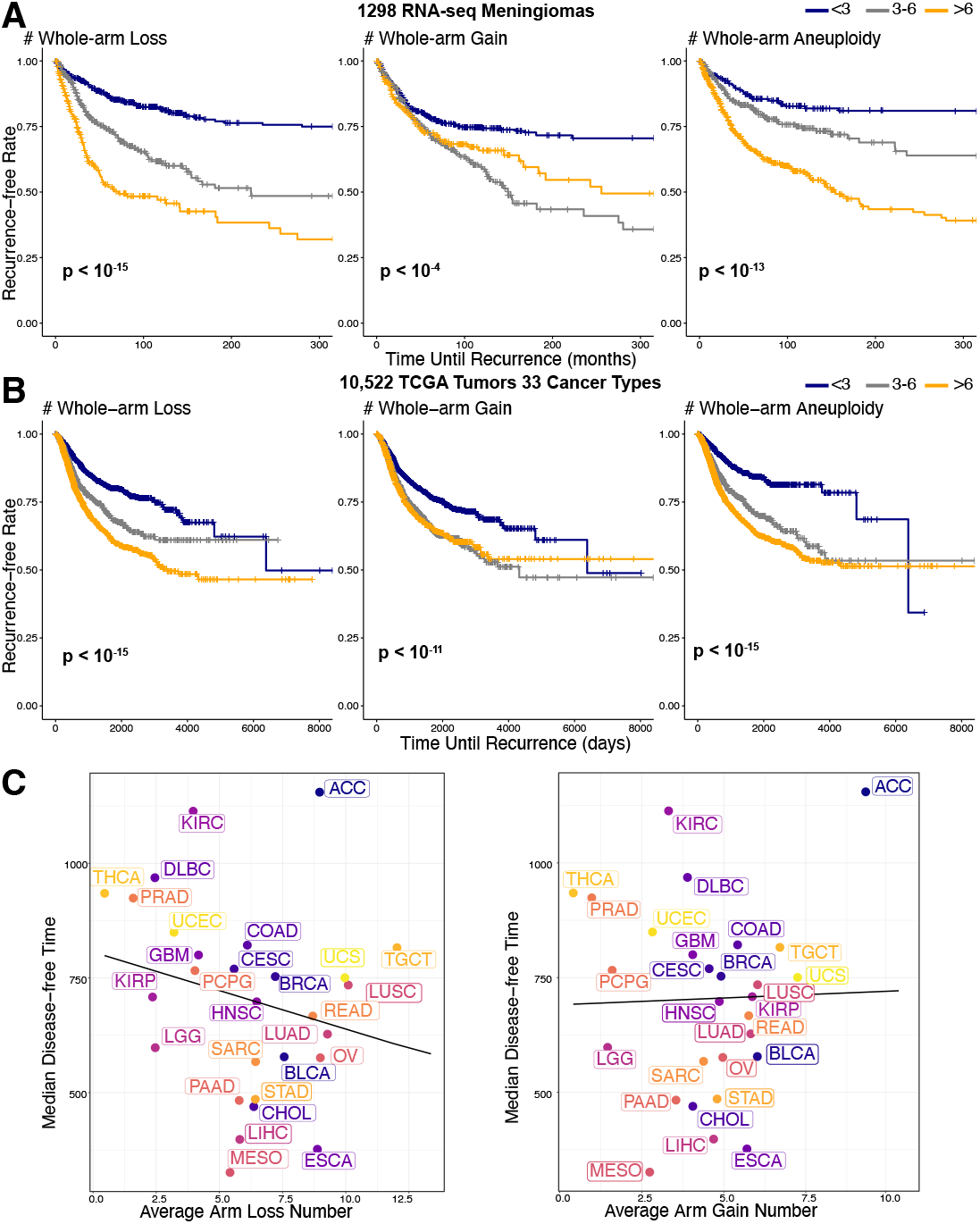
Whole-arm losses correlate with recurrence in meningioma and across 33 TCGA cancer types. (**A**) Whole chromosome arm gains and losses were identified using CaSpER across 1,298 meningioma RNA-seq samples. Whole-arm aneuploidy is defined as the total number of chromosome arm gains or losses. Kaplan-Meier curves compare recurrence times among patients with low (fewer than three), medium (three to six), and high (more than six) levels of chromosome arm alterations. The log-rank test p-values, shown in the lower-left corner of each panel, assess the statistical significance of survival curve separation. (**B**) Chromosome arm aneuploidy was determined using the ABSOLUTE algorithm in 10,522 TCGA whole-genome sequencing (WGS) samples, as reported in Ref. **(10)**, Table S2. Disease-free time serves as the recurrence metric. (**C**) Relationship between median disease-free time and the average number of whole chromosome arm losses (left) or gains (right) across 33 TCGA cancer types. The solid lines represent linear regression fits. The kidney chromophobe (KICH) cancer type has a significantly longer disease-free time, exceeding the range displayed in the figure.

The highly significant loss over gain bias in meningiomas that we observed in RNA-seq data encouraged us to ask whether the same bias can be detected in The Cancer Genome Atlas Project (TCGA) data. Examination of pan-cancer whole-genome sequencing (WGS) data from TCGA for 10,522 patients (10) revealed an overall average excess of whole-arm losses per patient (5.8) over gains (4.2) (Table S1).

To predict outcome based on whole-arm aneuploidies in TCGA pan-cancer data, we used recurrence (disease-free interval) data from a previous study (11), grouping by the same whole-arm aneuploid frequency intervals as for meningiomas. Strikingly, we observed qualitatively similar predictions for losses, gains and total aneuploids for pan-cancer data as for meningiomas, with the most rapid recurrence for >6 losses but not for >6 gains (Fig. 1B). This qualitative concordance between pan-cancer WGS data and meningioma RNA-seq data predictions of clinical outcome is especially notable considering the large variations in the average number of losses and gains between individual cancer types (Fig. 1C).

### Whole-arm are more frequent than whole-chromosome aneuploids across cancer types

Based on breakpoint analysis, whole-arm aneuploids account for 23 of the 25 most frequent breakage events and span on average 22.5% of the human genome in TCGA pan-cancer data (12). When whole-arm gains and losses are plotted as a fraction of the total for each cancer type, the overall frequencies are seen to vary over a wide range (Fig. 2 and Fig. S2). For example, adrenocortical carcinoma (ACC) averages 9.3 gains and 9.0 losses per patient, whereas acute myeloid leukemia (LAML) shows only 0.85 gains and 0.72 losses per patient. Arm-to-arm differences are also cancer type-specific, with very similar frequencies for all 39 autosomal arms for ACC but conspicuous arm-to-arm variations for glioblastoma (GBM). It is generally assumed that concordant p and q arm loss numbers in genomic data represent whole-chromosome losses.

**Figure 2.**
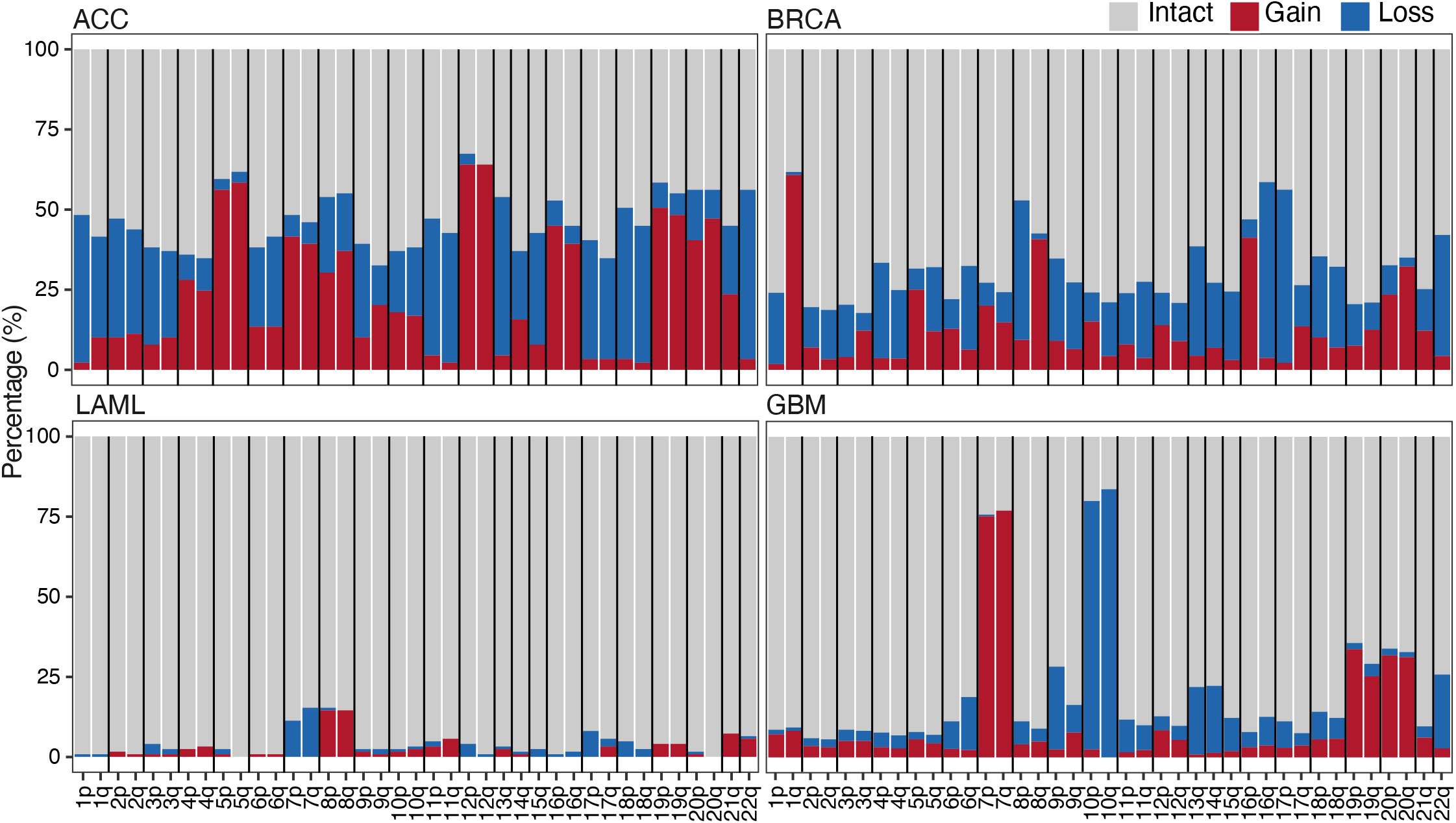
Percentage of whole-arm gains and losses in selected TCGA data. Each histogram bar represents the overall percentage of intact, gained and lost chromosome arms in TCGA data for the indicated cancer type (**Supplementary Table 1**), where ACC is adrenocortical carcinoma, BRCA is breast cancer, LAML is acute myeloid leukemia and GBM is glioblastoma. For clarity, only 4 of the 33 cancer types are shown. The full TCGA set of histograms is displayed in **Supplementary Figure 2**. Vertical black lines separate whole chromosomes, where Chromosomes 13, 14, 15, 21 and 22 are acrocentrics with a short p arm (not displayed) and long q arm.

Indeed, a molecular criterion for glioblastoma is gain of Chromosome 7 and loss of Chromosome 10 (13) with 7p and 7q gains and 10p and 10q losses in ∼80% of tumors compared with ∼10% total gains and losses for most other arms (Fig. 2). Based on concordance between p and q arms within a tumor, TCGA patient data revealed that overall 21.6% of the total are whole-chromosome aneuploids, and these show a small overall excess whole-chromosome losses over gains (1.4 losses versus 1.2 gains, (Table S1). Therefore, despite examples such as GBM, which is driven primarily by Chr7 gain and Chr10 loss (14), whole-arm gains and losses are by far more conspicuous across cancer types in TCGA.

### Metacentric and acrocentric aneuploidies occur at similar frequencies across cancer types

In cancer, breakpoints in centromeric and pericentric regions are on average 4.4 times more frequent than breakpoints in euchromatic arms based on breakpoint density along the chromosome (12). Because these regions consist of tandemly repetitive a-satellite DNA and the functional centromere accounts for only ∼5% of the total (15), the 4.4-fold excess of centromere-specific breaks is likely a gross under-estimate of the likelihood that a break in a functional centromere will result in a whole-arm aneuploid. As the vast majority of aneuploid chromosomes in cancer must have undergone centromere breakage, and this alone may result in failure to attach to the mitotic spindle, it is possible that aneuploidy occurs without any other mitotic error.

To determine whether mitotic errors, in addition to centromere breaks, are responsible for whole-arm gains and losses in cancer, we took advantage of the two classes of human chromosomes based on the position of the centromere. Metacentric chromosomes have two euchromatic arms (Fig. 3A) and so require a centromere break to generate a whole-arm aneuploidy (Fig. 3C). There are five human acrocentric chromosomes (13, 14, 15, 21 and 22), which have similar kinetochore conformations as metacentrics (16), but only a single euchromatic arm (Fig. 3B). Acrocentrics that are gained or lost by mitotic error will be effectively indistinguishable from those that have undergone a centromere break event (Fig. 3D). Acrocentric short arms comprise only redundant ribosomal DNA genes and other tandem repeats, and single short-arm gains or losses are not detected in genomic studies. Whole-chromosome meiotic or mitotic segregation errors occur, but centromere position does not predict frequency, based on pre-meiotic mitoses during human oocyte maturation (78% metacentrics expected, 83% observed, n = 52) (17). Thus, if mitotic errors are essential for the generation or perpetuation of a significant number of whole-arm aneuploids, then there should be an excess of acrocentrics (both those with and those without centromere breakpoints) relative to metacentrics (all of which must have centromere breakpoints).

**Figure 3.**
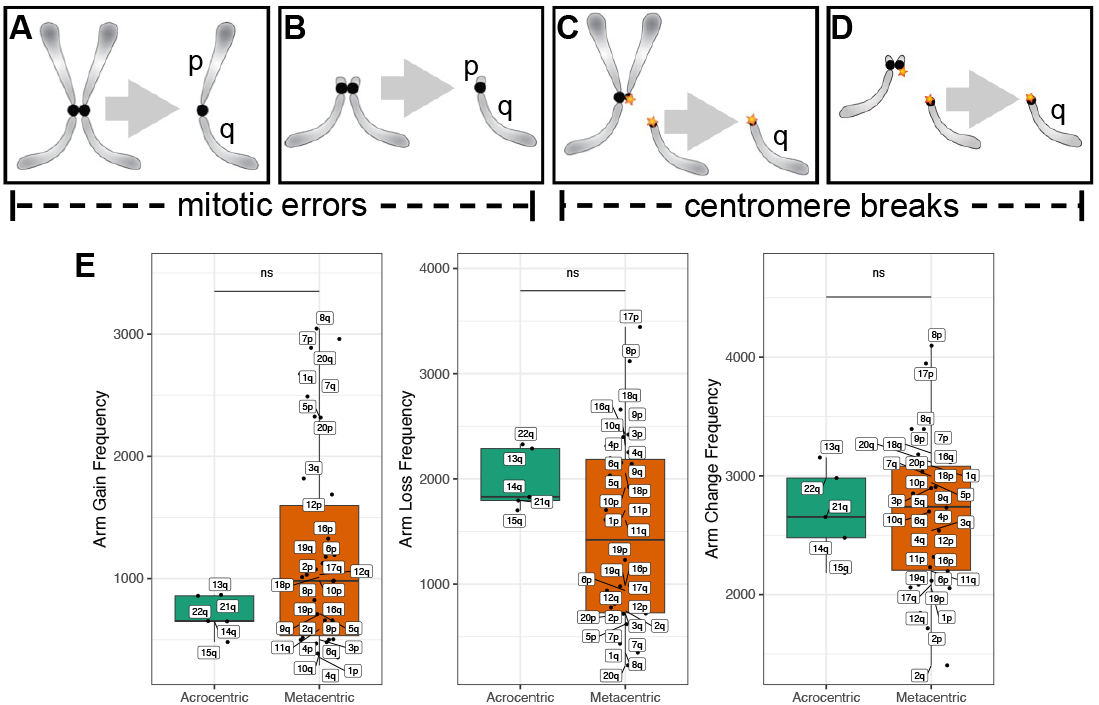
Acrocentrics and metacentrics gain or lose whole arms at similar frequencies in cancer. (**A**) A metacentric segregation; (**B**) An acrocentric segregation; (**C**) A metacentric segregation following a centromere break; (**D**) An acrocentric segregation following a centromere break. Acrocentric p chromosome arms are short and lack unique loci; thus intact acrocentrics are not distinguishable from q arms resulting from centromere breaks in genomic studies. (**E**) TCGA whole-genome sequencing (WGS) data by tumor type. Each dot represents a different autosomal chromosome arm (5 acrocentric long arms and 17 metacentrics).

To determine whether acrocentric are in excess over metacentric aneuploid chromosomes as expected for mitotic error in addition to centromere breaks, we have analyzed WGS data from TCGA for acrocentrics and metacentrics from 10,674 patients based on allele-specific copy number segmentation analysis across 33 cancer types. However, we observed no significant differences between acrocentrics and metacentrics in the frequency of whole-arm gains or losses or both (Fig. 3E and Fig. S3). We performed a similar analysis of long-read RNA-seq data from B-ALL and AML leukemias and again observed no significant differences (Fig. S4) (18). Considering that we are comparing all autosomal chromosome arms from >10,000 patient tumors, our inability to detect whole-arm events attributable to mitotic error is highly robust.

## Discussion

### Total whole-arm losses predict outcome across cancer types

In our previous study, we had found that RNAPII at the S-phase-dependent histone genes is increased in most human cancers, reminiscent of the observation that elevated histone expression promotes life span extension in yeast (21). We also showed that RNAPII at histone genes correlated strongly with total arm losses and speculated that histone overexpression results in centromere breaks that lead to aneuploidy (6). Here we have extended these findings by showing that simply counting the number of whole-arm losses in public RNA-seq and WGS data predicts clinical outcome in diverse human cancers better than whole-arm gains. This is counter-intuitive given that trisomies (3/2 gain) occur in ∼0.3% of newborns (22), but no autosomal monosomy (1/2 loss) is known to have ever come to term, and yet whole-arm losses are evidently more fit than gains in cancer (23, 24). As whole-arm aneuploids from metacentric chromosomes must be generated by centromere breaks, our finding that the degree of aneuploidy is patient-specific in general for all chromosome arms implies that whatever is causing centromere breaks is not chromosome-specific but rather is a general cellular event.

The relationship between aneuploidy and cancer has been vigorously debated ever since Boveri built on Hansemann’s careful observations with insights from genetics to argue that aneuploidy underlies the cancer phenotype (26). This debate has continued for over a century (27, 28), and recent evidence from TCGA data that whole-arm aneuploids result in either net gains of tumor drivers or losses of tumor suppressors (12) supports the Hansemann-Boveri hypothesis. In contrast, the question of whether mitotic errors cause aneuploidy has not been seriously challenged. Merotelic attachments of mouse chromosomes in cells with a Dido mutation were reported to break at anaphase (29), however, direct evidence for centromere breaks at anaphase is extremely limited. Furthermore, effective tension-dependent mechanisms have evolved to release attachments, sometimes resulting in lagging chromosomes (30). Therefore, in the absence of conclusive evidence that anaphase tension can break centromeric DNA of human chromosomes, we consider the generation of whole-arm aneuploids during mitosis to be unlikely to account for their high abundance in cancer.

### A single general cause of aneuploidies in cancer?

Our demonstration that centromere breaks alone can account for the large majority of aneuploidies in cancer focuses attention on the possibility that merotelic attachments to a single chromatid splits the centromere in two during anaphase (7). Centromeres are fragile sites in the genome (29, 31-34), but the amount of force required for breaking DNA (19) is vastly in excess of the 5-7 pN required to rupture a single kinetochore-DNA interaction (35). Although merotelic attachments of spindle microtubule bundles to a single kinetochore might in principle break centromeres at anaphase, chromatin is elastic (19), and single centromeres become distorted when pulled towards opposite poles, perhaps owing to the bipartite organization of the centromere (16, 20).

The patient-specific whole-arm loss bias that we observed across tumor types and in RNAPII, RNA-seq and WGS data points to a general cause of aneuploidy in cancer that similarly affects all chromosome arms in each patient. This possibility was envisioned by Boveri, who proposed that an “abnormal event” in the primordial cancer cell precedes aneuploidy (26). To explain the generation of whole-arm aneuploids, we have proposed that excess H3 histones resulting from overexpression of S-phase-dependent histone genes in cancer compete with CENP-A for incorporation at centromeres (6) (Fig. 4A). CENP-A is the histone H3 variant that marks active centromeres and is essential for kinetochore function (16). Depletion of CENP-A during S phase can lead to transcription-replication conflicts and R-loops that stall the replication machinery, resulting in fork collapse and chromosome breaks of the type that can lead to whole-arm aneuploids (36-38) and replacement with H3 nucleosomes (39, 40). As these S-phase events occur during a different phase of the cell cycle than mitotic errors, S-phase and mitosis models are mutually exclusive in that a centromere break could have occurred before a mitotic error or vice versa, but not at the same time. However, it is possible that mitotic segregation errors help to perpetuate whole-arm aneuploids once they are generated by a break. Although 21.6% of aneuploidies were scored as whole-chromosome aneuploids in TCGA data (10), breaks between functionally bipartite human centromeres with merotelic attachments to each half-centromere (16) would result in separated p and q arms, and simultaneous gain or loss might be mis-scored as whole-chromosome aneuploidies.

**Figure 4.**
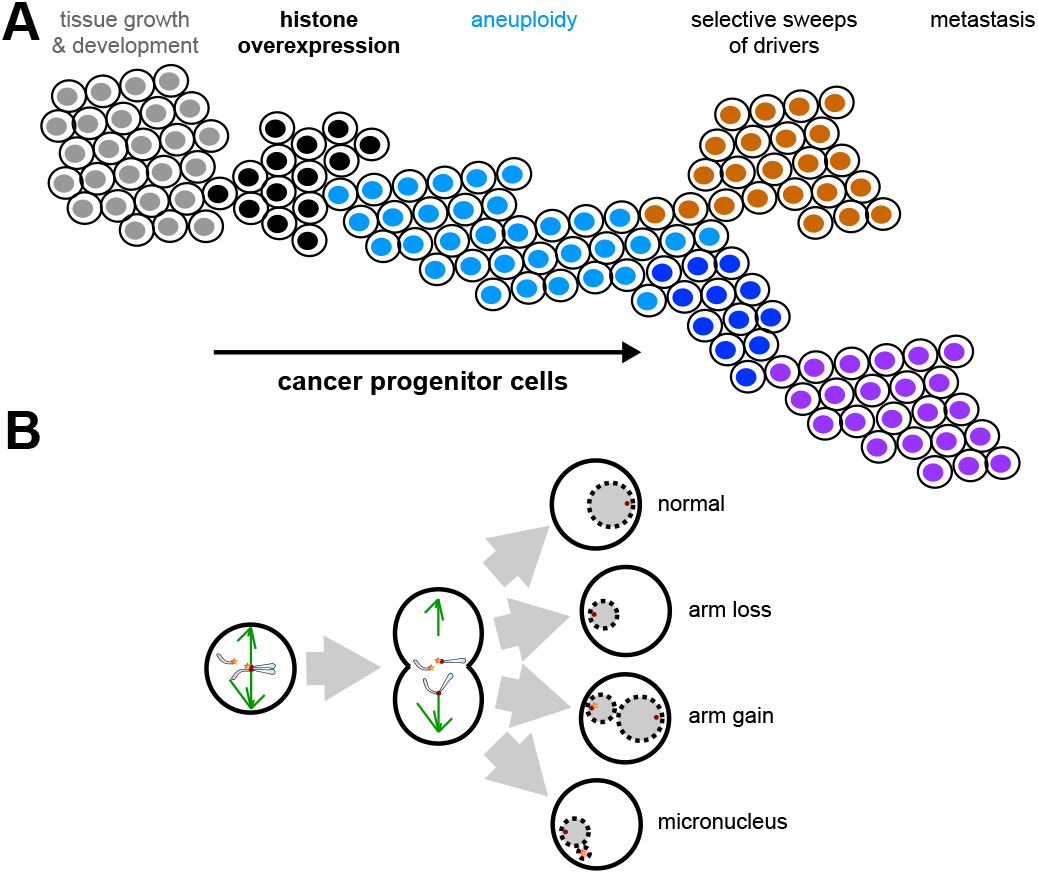
Model for generation of centromere breaks leading to losses > gains. (**A**) Cancer progression: Normal cells (grey) proliferate and differentiate to populate tissues, but aberrant induction of histone overexpression (black) promotes both hyperplasia and chromosome instability. Selection in hyperproliferating clones drives the frequencies of certain chromosomal abnormalities and the evolution of malignant cellular features (brown, dark blue, purple). (**B**) Model: Defective centromeres compromised by centromeric histone displacement will break, leading to widespread aneuploidy (blue) through whole chromosome arm loss, arm gain, and through whole chromosome loss. The occurrence of micronuclei by encapsulation of fragmented chromosome arms further stimulates chromosomal instability.

How might centromere breaks account for the excess of whole-arm losses over gains? Hypertranscription and over-expression of S-phase-dependent histones compete with CENP-A nucleosomes (40), resulting in centromere breaks (Fig. 4B). If a centromere break releases a chromosome arm that lags at anaphase, it may form a micronucleus upon mitotic exit (41), eventually to undergo degradation in the cytoplasm resulting in a whole-arm loss. However, in cases where S-phase replication occurs within a micronucleus and is followed by kinetochore formation on the broken centromere and monopolar microtubule capture, then the nucleus reforming at that pole during the following telophase will include two extra chromosome arms. In this way, whole-arm gains would be secondary events, sometimes leading to replication but at other times to degradation and so would be less frequent than whole-arm losses, as we observed.

As whole-arm gains and losses are by far the dominant aneuploid type in cancer (12), prevention of centromere breaks could be a general therapeutic strategy. We suggest that preventing over-expression of histone genes might be such a strategy. S-phase-dependent histone genes have unique therapeutic vulnerabilities. These genes are present in clusters located in phase-separated histone locus bodies (HLBs) where they are the only RNAPII transcripts with 3’ stem-loops that are recognized by the stem-loop binding protein (42). S-phase-dependent 3’-end RNA processing within the HLB is mediated by the U7 small nuclear ribonucleoprotein complex without polyadenylation. Also, S-phase-dependent histone genes are uniquely activated for transcription by the NPAT protein, which is the major structural component of the HLB, and are uniquely repressed outside of S-phase by soluble histone H4 (43). Like RNAPII, which is the target of multiple general anti-cancer therapies, the S-phase-dependent system of gene regulation is ancestral for eukaryotes (44), and so anti-histone therapies may be less likely than targeted therapies to induce resistance. Thus the generality of the whole-arm loss bias that we have shown may open the way to development of entirely new general anti-cancer therapeutic options.

## Materials and Methods

### RNA-seq data processing and analysis

The RNA-seq data for 1,298 meningioma samples and corresponding recurrence clinical data were obtained from 13 meningioma studies compiled by Thirimanne et al. (8). Raw FASTQ files w ere r e-aligned using STAR (version 2.7.11) to the hg19 reference genome from GENCODE (v19). Unstranded RNA-seq counts were used as raw counts per gene per sample.

Chromosome arm gains and losses were inferred using CaSpER (9), with the function extractLargeScaleEvents from the CaSpER R package. The default threshold of 0.75 was applied for calling large-scale chromosomal alterations. Kaplan-Meier survival curves were generated using the survminer R package, and log-rank tests were performed to evaluate the significance of the differences in survival curve distributions.

### Whole-genome sequencing data analysis

Whole-genome sequencing (WGS) data and associated survival information were obtained from The Cancer Genome Atlas (TCGA) (https://portal.gdc.cancer.gov). Whole chromosome arm aneuploidy was determined using the ABSOLUTE algorithm (45), and results were directly retrieved from Table S2 of Taylor et al. (10).

To assess recurrence outcomes, we used disease-free interval (DFI) as the primary survival metric, as recommended (11). Statistical comparisons of whole chromosome arm aneuploidy between acrocentric and metacentric chromosomes were conducted using the Wilcoxon rank-sum test, with significance levels indicated by corresponding p-values in boxplot analyses.

## Supporting information

Peer Reviewers

## Acknowledgments

We thank members of the Henikoff laboratory, Sue Biggins for insightful discussions and both Peer Reviewers. S.H. is a Howard Hughes Medical Institute Investigator. This work was supported by National Institutes of Health grant HG012797 to Y.Z.

## Author Contributions

Y.Z, K.A. and S.H. designed research; Y.Z., K.A. and S.H. performed research; Y.Z., K.A. and S.H. analyzed data; and Y.Z., K.A. and S.H. wrote the paper.

---

**Fig. S1.**
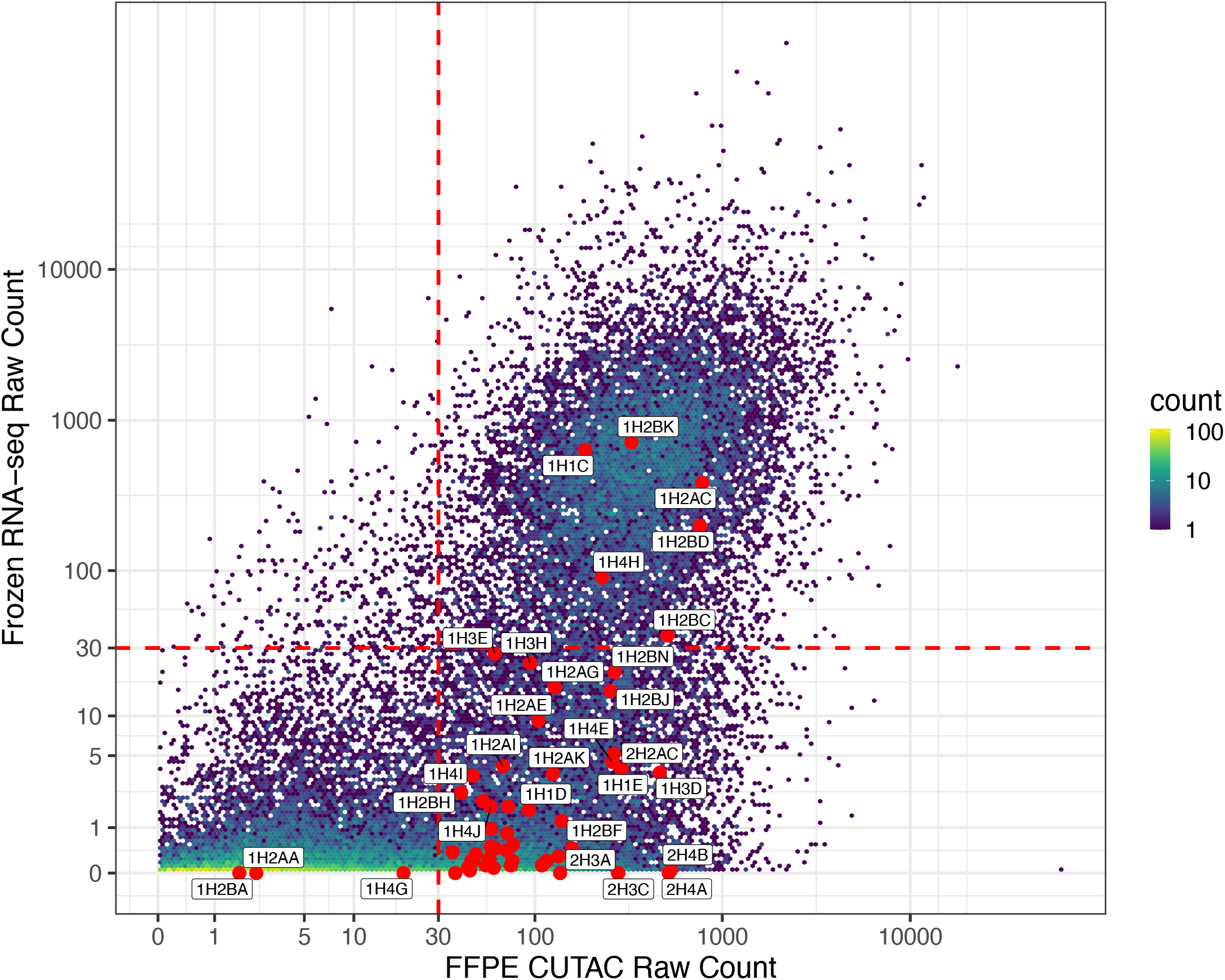
S-phase-dependent histone mRNAs are under-represented in meningioma patient RNA-seq data. Log-scale hexbin plot compares the gene-by-gene distribution of average RNAPII fragment counts from FFPE-CUTAC to average RNA-seq transcript counts from frozen meningioma samples of the same patients. The individual histone gene signals are indicated as red circles, where the large majority are high in RNAPII but very low for RNA-seq. Exceptions in the upper right quadrant are likely to represent S-phase-independent replacement histones, including histone partners for H2A.X, H2A.Z, H3.3 and CENP-A variant histones, which are encoded by intron-containing genes outside of the S-phase-dependent histone clusters.

**Fig. S2.**
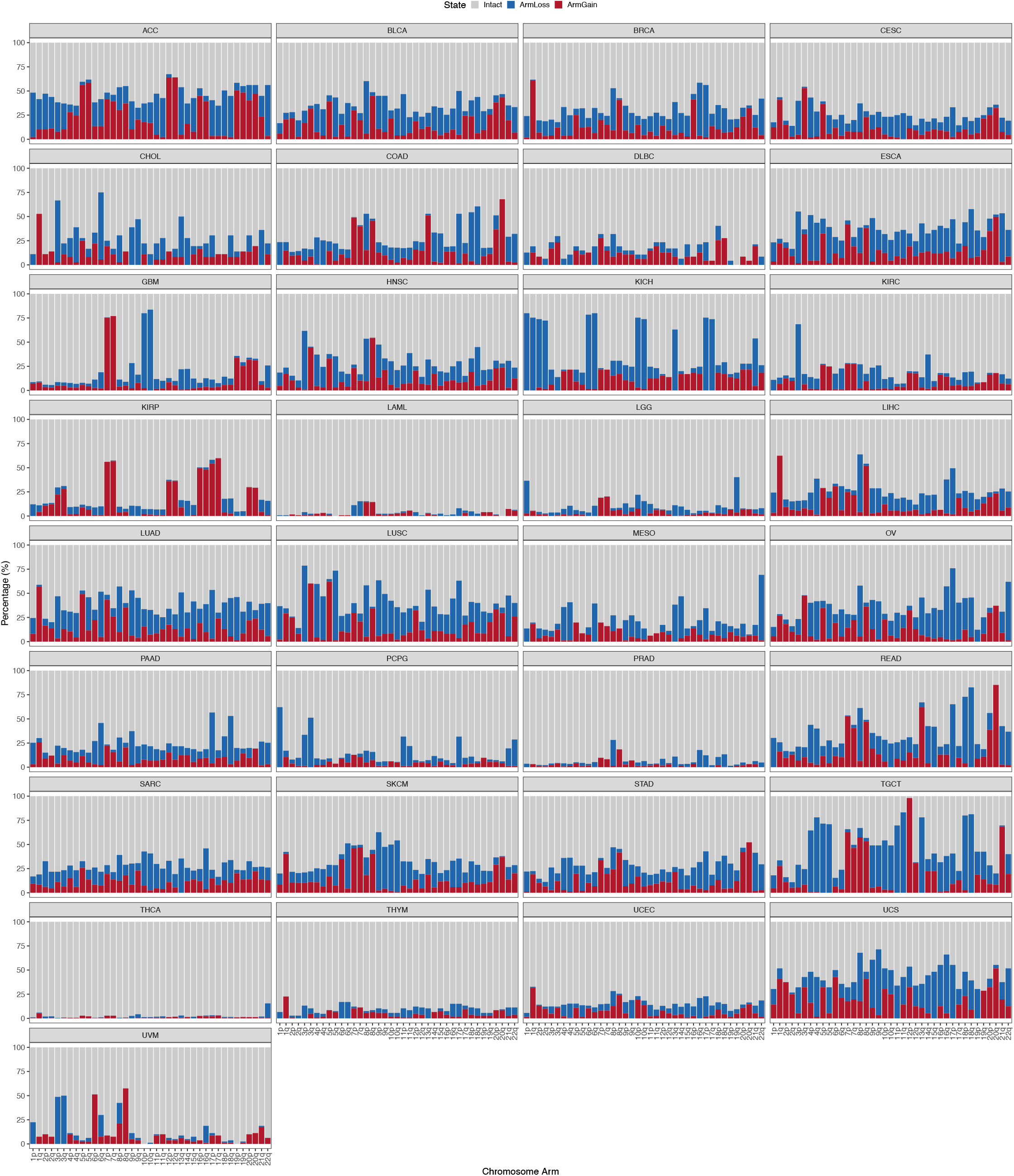
Percentage of whole-arm gains and losses in TCGA data. See the legend in Figure 2.

**Fig. S3:**
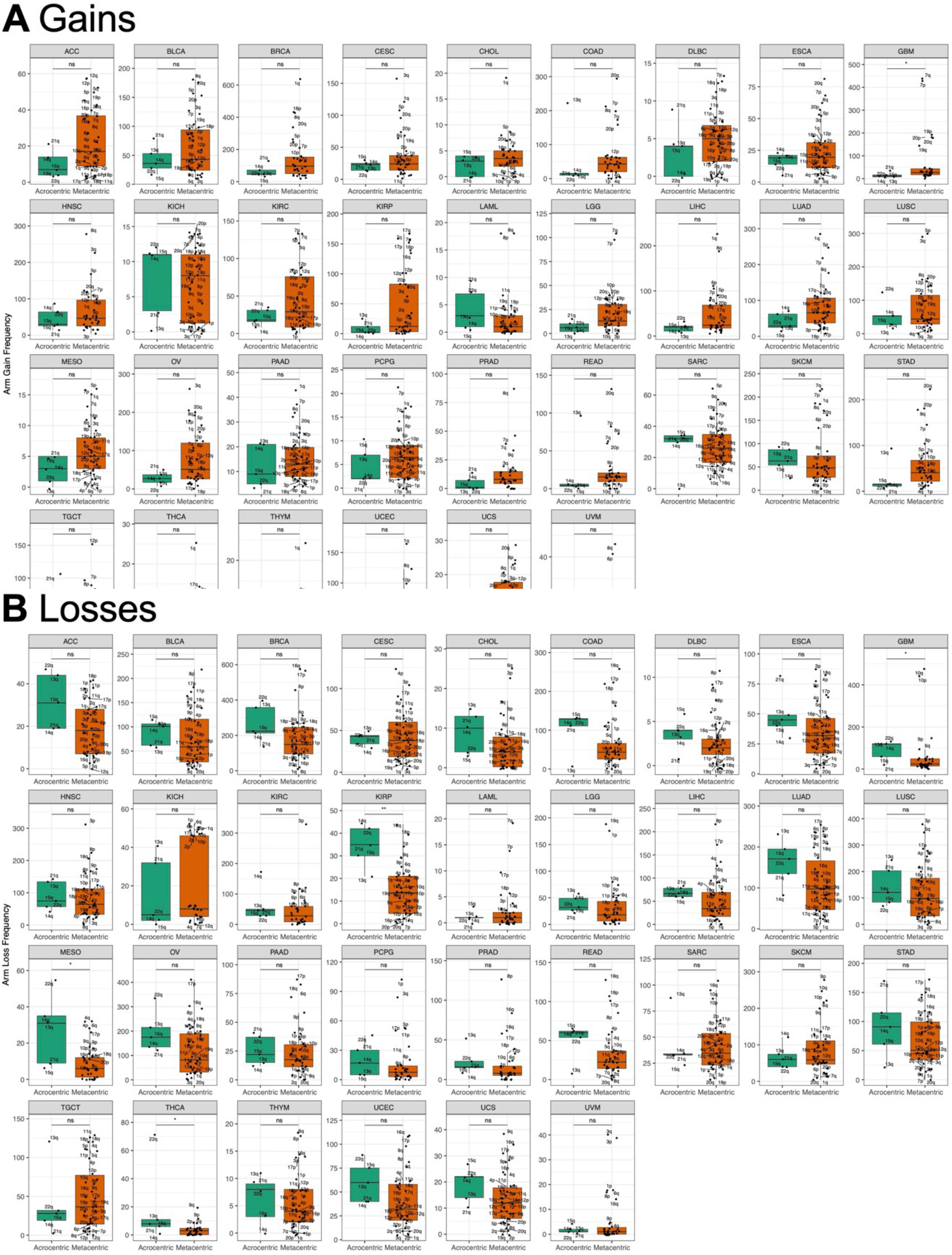
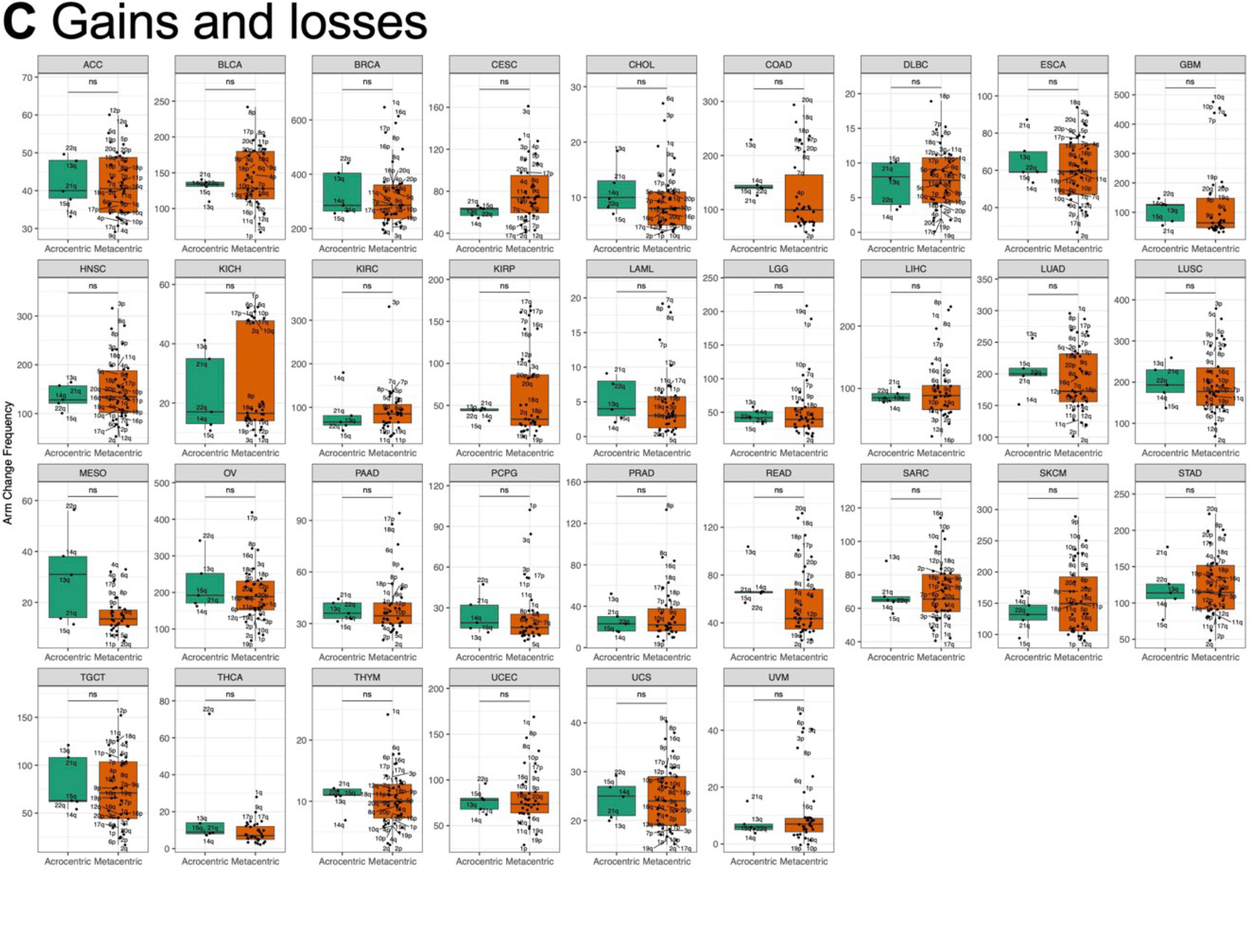
Acrocentric and metacentric whole-arm aneuploidies are recovered at similar frequencies in 33 cancer types. We used whole-genome sequencing data from 10,674 cancer patients spanning 33 cancer types downloaded from The Cancer Genome Atlas (TCGA, https://portal.gdc.cancer.gov). For 32 of 33 cancer types, no significant differences are seen between acrocentrics and metacentrics in the frequencies of whole-arm gains, losses or both gains and losses (Wilcoxon rank test), except for OV, in which metacentrics are in excess at p < 0.05. Each dot represents a different autosomal chromosome arm (5 acrocentric long arms and 17 metacentrics). We inferred the chromosome arm gain or loss using ABSOLUTE (38) profiles for each patient. We summed the major and minor alleles of each segment and took the minimum and maximum of the copy number across segments on each chromosome arm. Any increases in the minimal allelic copy number from the diploid copy number of 2 indicate an arm gain. Similarly, any decreases in the maximum allelic copy number from the diploid copy number of 2 indicate a whole-arm loss for the corresponding autosomal arm. (**A**) Gains; (**B**) Losses; (**C**) Gains or losses.

**Fig. S4:**
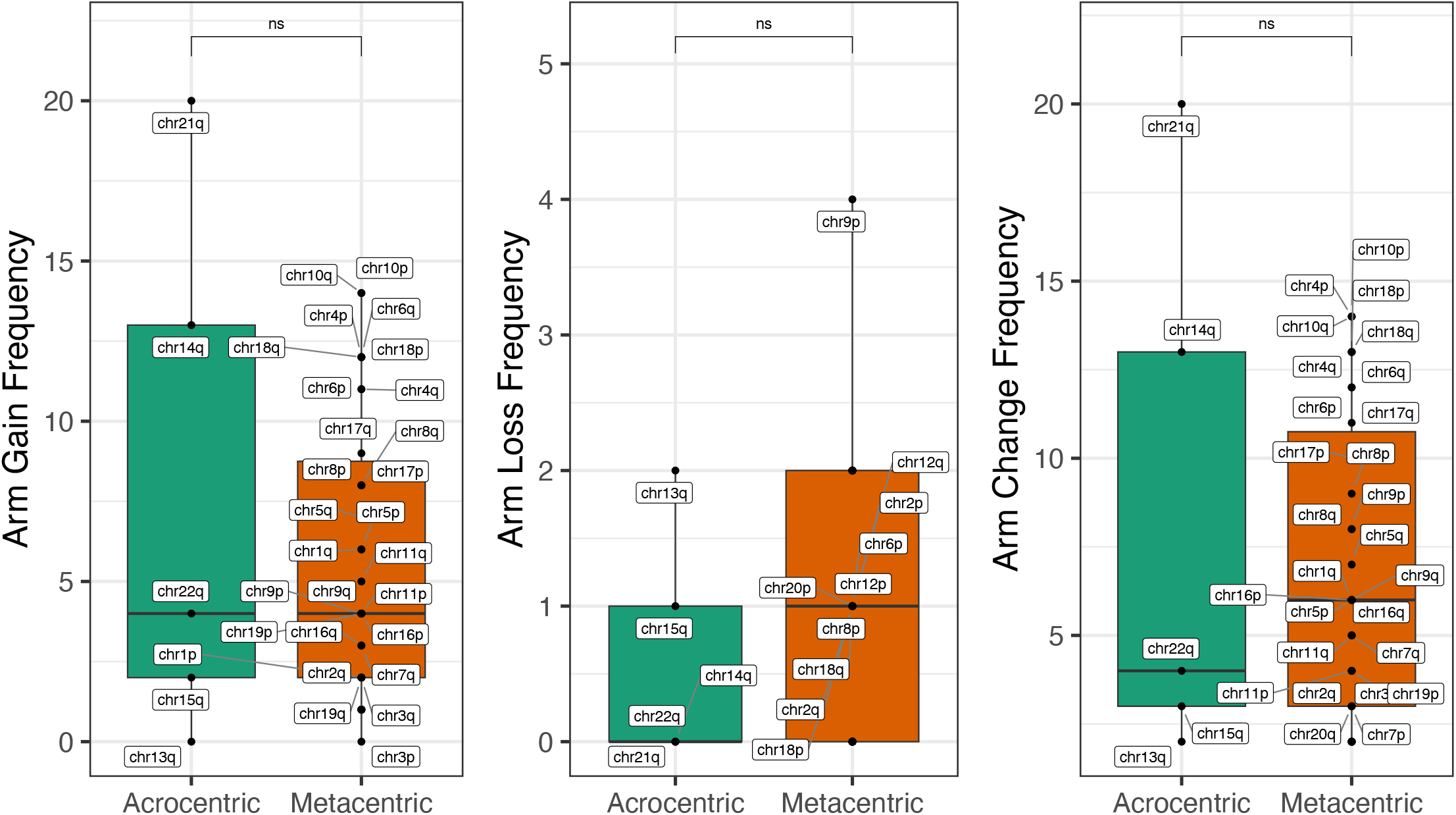
Acrocentric and metacentric whole-arm aneuploidies are recovered at similar frequencies in Nanopore long-read RNA-seq data. See the legend in Figure 3. Data are from combined B-ALL and AML leukemia patient samples described in Ref (18).

