## Supplementary material for "Total whole-arm chromosome losses predict malignancy in human cancer": Peer Reviewers

### Peer Review File

#### Reviewer 1

This is an interesting and timely article detailing the potential initiating event in aneuploidies and propensity for cancerous outcomes. The paper follows up an initial finding that RNA polymerase II at histone genes is correlated with whole-arm chromosomal aneuploidies. The authors conclude that the initiating events are centromere breaks. This is important for the field and consistent with recent findings of centromere fragility (doi: 10.1016/j.molcel.2024.01.018; doi: 10.1016/j.dnarep.2023.103569; doi: 10.1038/s41467-025-57277-9).

**We have added the additional references on centromere fragility in the new opening paragraph of the Discussion section: "A single general cause of aneuploidies in cancer?".**

The authors begin the results citing a paper that purports to demonstrate aneuploidy is generated from centromeric or pericentromeric regions. The paper is a review and propagates a common misperception that spindle forces are strong enough to break double-stranded DNA. In Alonso-Guerrero et al., http://www.pnas.org/cgi/doi/10.1073/pnas.0912143106, the authors present primary data and develop the spindle breakage argument. Alonso-Guerrero uses number for microtubule forces about 10X higher than shown in authentic kinetochores. A-G cites pioneering work from C. Rieder, that was focused on lateral chromosome-microtubule interactions, rather than kinetochore microtubules. More recent studies from the Asbury lab demonstrate forces on the order of 5-7 pN for single microtubules. A detailed commentary on forces required for chromosome breakage can be found in citation listed in ref 18.

**We had mentioned the problems with the van Wely model in the Discussion section, but for the Introduction, we cited the van Wely review because after 120 years of aneuploidy in cancer, it was the first to our knowledge to propose a single underlying mechanism for aneuploidy and centromere breaks in cancer as opposed to a collection of unrelated mechanisms. To give the authors credit for raising this issue, we have modified the introductory statement to avoid the implication that we subscribe to the authors' model for breakage: "Whole-arm losses and gains of metacentric chromosomes are generated by breaks in centromeres or pericentromeric regions*, although the underlying mechanisms have been speculative* [7]."**

**The points made by Reviewer 1 are more appropriately addressed in the Discussion section "A single general cause of aneuploidies in cancer?" where we now start out with a paragraph including the evidence cited by the Reviewer critical of the merotelic attachment breakage model highlighted in the van Wely review, including the additional points made in the original reference 18 (now reference 32) and the additional evidence for centromere fragility referred to above.**

The authors state that the functional centrumere is only 5% of the a-satellite (line 155). They cite a Biorxiv paper Logsdon et al.,. I couldn't find the basis for the citation in

reference 15. I could only find that there were two hypomethylated regions for 7% of the centromeres. The authors should clarify this statement in the results.
**This was an Endnote error, now fixed. The ~5% value for the functional centromere is from this paper from the T2T project: G. A. Logsdon et al., The variation and evolution of complete human centromeres Nature 629, 136-145 (2024).**

The authors state on line 176 "To distinguish these two models...." I couldn't determine which models they were distinguishing from one another. Maybe between breaks of metacentrics vs non-disjunction of acrocentrics? The authors should clarify.
**Now clarified: "*To determine whether acrocentric are in excess over metacentric aneuploid chromosomes as expected for mitotic error in addition to centromere breaks, we have analyzed....*"**

Finally, the authors state that "Hypertranscription and ... compete with CENP-A nucleosomes" citing Henikoff, 2025. There is no data in Henikoff 2025 for this statement. Instead Henikoff 2025 reads "To explain how overproduction of histones might account for this notable whole-arm loss bias in patient tumors, we propose that excess H3 histones compete with CENP-A histones at S-phase for nucleosome assembly at centromeres" and cite other papers. The authors clarify what is proposed from what and who reported the data.
**That was an erroneous citation now corrected. We should have cited Ref. 32 (Blower et al. 2002), which is now reference 40, which showed that excess H3 produced in Drosophila cells causes loss of CID/CENP-A.**

**<u>Reviewer 2</u>**
In this study Zheng et.al test the hypothesis of centromere breaks as a driver of aneuploidy and whether whole arm aneuploids can have predictive value of cancer outcomes utilizing large RNA sequencing and whole genome sequencing datasets. They also perform a creative analysis, comparing the rates of chromosome arm aneuploidies in acrocentric and metacentric chromosomes, to assess the relative importance of mitotic errors vs. centromere breakages in generating/perpetuating whole arm gains and losses. Their approach is computational and is potentially a novel approach to studying an old observation studied in the context of Cancer Biology.
Specific comments:
1. Based on their recent publication in Science, the authors implicate inappropriate RNA Pol II targeting and potential competition with histone H3 variant CENP-A in centromere breaks and whole arm losses. How does histone overexpression data correlation look when comparing 10522 TCGA tumours across 33 cancer types?
**There is no S-phase-dependent histone mRNA data for TCGA samples. This is because all histone mRNAs are 3'-end processed by U7 snURP and lack poly(A) tails, making them essentially invisible in RNA-seq studies, which rely on poly(A) priming. To avoid confusion on this point we have added a sentence to the end of the first paragraph of the Results section: "*S-phase-dependent histone mRNAs are the only RNAPII-transcribed protein-coding genes that are not 3'-polyadenylated, and so they are essentially absent from RNA sequencing datasets (Supplementary Figure 1).*"
The new Supplementary Figure 1 compares the gene-by-gene distribution of**

**average RNAPII fragment counts to average RNA-seq transcript counts from frozen meningioma samples. It shows that the large majority are high in RNAPII but very low for RNA-seq. Exceptions are likely to represent S-phase-independent replacement histones, including histone partners for H2A.X, H2A.Z, H3.3 and CENP-A variant histones, which are encoded by intron-containing genes outside of the S-phase-dependent histone clusters.**

2. Fig 2 will be more informative and will have broader scopes if additional cancer subtype data is added on.
**We only showed the 4 panels referred to in the text. A problem with adding more panels is that the choices would be arbitrary and/or redundant, so we do not agree that this would make the figure more informative or meaningfully expand the scope, which is why we place the full set of 39 autosomal whole-arm histograms in the supplement.**

3. In line 154, the authors begin with "Because these regions". Some clarification of where the regions are coming from will be very helpful for the readers.
**To avoid ambiguity, we have now reworded the previous sentence to: "In cancer, breakpoints in centromeric and pericentric regions are on average 4.4 times more frequent than breakpoints in euchromatic arms based on the per-megabase density of breakpoints [12]. Because these regions..."**

4. The argument put forth on lines 160-175 is intriguing. Figures 3A-D, which help to explain and visually support the argument, may benefit from a few additions for clarity: for instance, by highlighting the "invisibility" of the short arm of acrocentric chromosomes, and also by labelling on the figure panels which situations are arising from a "mitotic error".
**We have enhanced the Figure 3A-D cartoons as suggested by Reviewer 2.**

5. Though it may be beyond the scope of the current study, is it possible to examine arm-specific aneuploidies of acrocentric chromosomes from long-read sequencing?
**Although there are no long-read sequencing datasets in TCGA, there is a recently published study with available Nanopore sequencing data for B-ALL and AML leukemias (new Reference 18). We performed the same analysis as for the TCGA data and again there is no significant difference between metacentrics and acrocentrics in the frequency of whole-arm aneuploids. We have added these new analyses as Supplementary Figure 4.**
